# Locus-specific chromatin proteomics using dCas-guided proximity labelling in *Aspergillus nidulans*

**DOI:** 10.1101/2024.10.15.618449

**Authors:** Thomas Svoboda, Dominik Niederdöckl-Loibl, Andreas Schüller, Karin Hummel, Sarah Schlosser, Ebrahim Razzazi-Fazeli, Joseph Strauss

## Abstract

Proximity labelling that uses promiscuous biotin ligases (BirA) fused to a bait protein is a powerful tool to identify protein interaction partners *in vivo* under different metabolic or developmental conditions. BirA can also be used to determine protein composition and interaction partners at specific chromatin locations when it is fused with enzymatically-disabled Cas9 (dCas9) and then guided to the location of interest by sgRNAs. We adapted this method (called CasID) for fungal cells using the nitrate assimilation gene cluster of *A. nidulans* as a model locus and estrogen-inducible expression of the dCas9-BirA fusion to improve condition-specific labelling. For method establishment, we first verified the presence of dCas-BirA and a known transcription factor at the nitrate locus by chromatin immunoprecipitation (ChIP). Results show that both dCas-BirA and the AreA transcription factor are present at the locus of interest under the conditions used for biotinylation. We then optimized the CasID procedure for efficient labelling and background reduction using the CasID-sgRNA strain and two control strains, one lacking the sgRNA and another one lacking the whole CasID system. Here we provide proof-of-concept for the suitability of the method by showing that biotinylated proteins are enriched in the CasID strains in comparison to the controls. After background reduction, 32 proteins remained in two independent experiments exclusively enriched in the Cas-ID-sgRNA strain. Among these proteins was NmrA, an AreA-interacting regulator, and we also found several chromatin-associated proteins. Overall, our results demonstrate that Cas-ID is suitable for locus-specific labelling and identification of chromatin-associated proteins and transcription factors in *A. nidulans*. However, the high background of proteins that are biotinylated out of chromatin context or unspecifically attach to the affinity purification matrix needs to be addressed by implementing a set of rigorous controls. In summary, we herewith provide a detailed protocol for application of the method that proved to be useful for the identification of novel chromatin-associated proteins and their interaction partners at a specific genomic locus in divers metabolic and developmental conditions.

**Author summary:** This study demonstrates that locus-specific proteomics can be carried out by dCas-BirA guided proximity labelling in *Aspergillus nidulans.* For establishment, we targeted the well-described bidirectional promoter region between *niaD*, a nitrate reductase, and *niiA*, a nitrite reductase. At this locus we could test by chromatin immunoprecipitation (ChIP) in combination with qPCR if both, the dCas9-BirA fusion as well as a central transcription factor are at the locus under the conditions of our Cas-ID experiment. After this first control step, we considered that unspecific labelling by dCas-BirA during the time from translation to landing at the targeted chromatin locus may be one of the most relevant drawbacks of the method. Therefore, we developed a number of control strains that would allow us to clearly discriminate between background and sgRNA-dependent specific labelling at the locus. Our protein MS results validated these estimates and only considering the results of these controls enabled us to distinguish the set of locus-specific proteins from a very high general background. Finally, enrichment of biotinylated proteins through affinity purification with streptavidin resin and subsequent LC-MS/MS analysis showed that more than 800 proteins were detected in each sample, emphasizing the high background of the purification method. After background reduction of the control samples, we were able to identify 32 proteins which were exclusively detected in the test strain in two independent measurements, including several chromatin-associated proteins and NmrA, a negative regulator of the nitrate locus transcription factor AreA.

## Introduction

Proximity-dependent labelling is a molecular tool to assess interaction partners of proteins in living cells and is frequently used as an alternative to co-immunoprecipitation (co-IP) or other affinity-tag based enrichment techniques for protein complexes. The advantage of proximity labelling over the other techniques is that it also detects weak or only transient interactions and the method has been developed and optimized over the last decade (reviewed by Samavarchi-Tehrani et al., 2020). The most commonly used labelling molecule is biotin and there are biotin ligases as well as peroxidases available for biotinylation approaches. While HRP or APEX require biotin-phenol as well as peroxide as substrate and attach it to tyrosine, the promiscuous biotin ligase BirA (or an engineered version of it named TurboID), uses biotin as substrate and attaches the activated biotinyl-5’-AMP to lysine residues. Also different other versions like mini TurboID are available (reviewed by Samavarchi-Tehrani et al., 2020). Proximity labelling employing BirA or derivatives of it has already been used in many different organisms ranging from mammalian cells (Cho et al., 2020), Zebrafish (Xiong et al., 2021), different plant species (Yang et al., 2021) to yeast (Singer-Krüger and Jansen, 2022) and recently also to *Aspergillus nidulans* (Georgiou et al., 2023).

Beside fusing the biotin ligase to a bait-protein for which the interaction partner(s) should be identified, also fusion to enzymatically disabled “dead” Cas9 (dCas) is possible. dCas is still able to interact with single guide RNAs (sgRNAs) and bind the target location in DNA, however, due to a specific mutation, it cannot cut the DNA anymore. This allows the construction of a fused dCas-BirA chimera that produces a biotinyl-5’-AMP cloud at the target chromosomal location thereby biotinylating proteins that are in close proximity within a radius of approximately 30nm. dCas can be guided to a defined locus at the DNA by a designed sgRNA. This method thus allows locus-specific proteomics through the guidance of the proximity labelling system to a specific genomic locus in the living cell and to assess there the chromatin composition under defined experimental conditions (reviewed by Anton et al., 2018).

In this study we established the CasID protocol expressing TurboID-dCas and sgRNAs in *Aspergillus nidulans* which genome consists of 8 chromosomes encoding approximately 12.000 genes. One of the first metabolic pathways that were studied on a detailed genetic and molecular level using this model fungus was nitrate assimilation. In bacteria, fungi and plants this pathway allows to use oxidized nitrate as nitrogen source for growth and to reduce it to ammonium for the further incorporation into amino acids (Cove, 1979). A wealth of knowledge is available on the *cis-*acting regions and *trans-*acting factors regulating transcription of the genes involved in nitrate assimilation (Burger et al., 1991; Gallmetzer et al., 2015; Muro-Pastor et al., 1999; Punt et al., 1995; Scazzocchio, 2000; Scazzocchio and Arst, 1989; Schinko et al., 2010; Strauss et al., 1998). The genes are induced by the nitrate-specific C_6_Zn_2_-family transcription factor NirA that interacts with the GATA-type transcriptional activator AreA and together they synergistically activate their target genes (Berger et al., 2008; Berger et al., 2006; Kudla et al., 1990). Under nitrate-inducing conditions, NirA switches conformation, accumulates in the nucleus and both regulators bind to the promoter regions of nitrate-responsive genes (Bernreiter et al., 2007; Gallmetzer et al., 2015). AreA recruits histone acetylation to chromatin (Muro-Pastor et al., 1999; Muro-Pastor et al., 2004) and activates transcription synergistically with NirA. To prevent waste of energy, AreA is inactivated when the intracellular levels of already reduced nitrogen sources such as ammonium or glutamine are already high. Previous binding experiments using *in vivo* footprinting (Muro-Pastor et al., 1999; Muro-Pastor et al., 2004) or ChIP of an HA-AreA strain (Berger et al., 2008; Berger et al., 2006) demonstrated that binding of AreA is lost upon repression by ammonium. The mechanism behind AreA inactivation has been studied in detail and was found to be modulated by NmrA, a regulatory protein able to monitor the nitrogen status of the cell and eventually block AreA activity when reduced nitrogen levels are high (Lamb et al., 2003; Langdon et al., 1995; Stammers et al., 2001; Wong et al., 2007). Because these molecular genetic details of the transcriptional regulation and associated factors are available, we used the bidirectional promoter region of the divergently transcribed nitrate reductase (*niaD*) and nitrite reductae (*niiA*) genes as a model for the establishment of CasID in this fungus. In this study we describe the establishment and the results of locus-specific proteomics at the described *niiA-niaD* bidirectional promoter (from here on termed “intergenic region”, or “IGR”) using the dCas9-guided proximity labelling approach and appropriate control strains.

## Material and Methods

### Strains and growth conditions

#380 (additional wild type strain, no tag): *Aspergillus nidulans* wild type strain without any modification used as a negative control for all other strains. Genotype: *yA2; biA1; veA1*

#871 (HA-AreA): *Aspergillus nidulans* strain with the transcription factor AreA tagged with 3xHA. Genotype: *HA-AreA, pyroA1 or pyroA4, argB2, biA1, nku::bar, wA1*

#880 (CasID −sgRNA): in the strain #871, the TurboID-dCas fusion protein construct was introduced, but not the sgRNA expressing construct. Genotype: *HA-AreA, pyroA1 or pyroA4, argB2::pTurboID-dCas9, biA1, nku::bar, wA1*

#881 (CasID +sgRNA): Based on strain #880 that already expressed the TurboID-dCas fusion protein, a constract expressing an sgRNA that targets the TurboID-dCas fusion protein to the *niiA-niaD* promoter region next to AreA binding site 4 (see Fig 1). Genotype: *HA-areA, pyroA1 or pyroA4, argB2::pTurboID-dCas9&revsgRNA-nirAII/areAIV, biA1, nku::bar, wA1*

**Figure 1:**
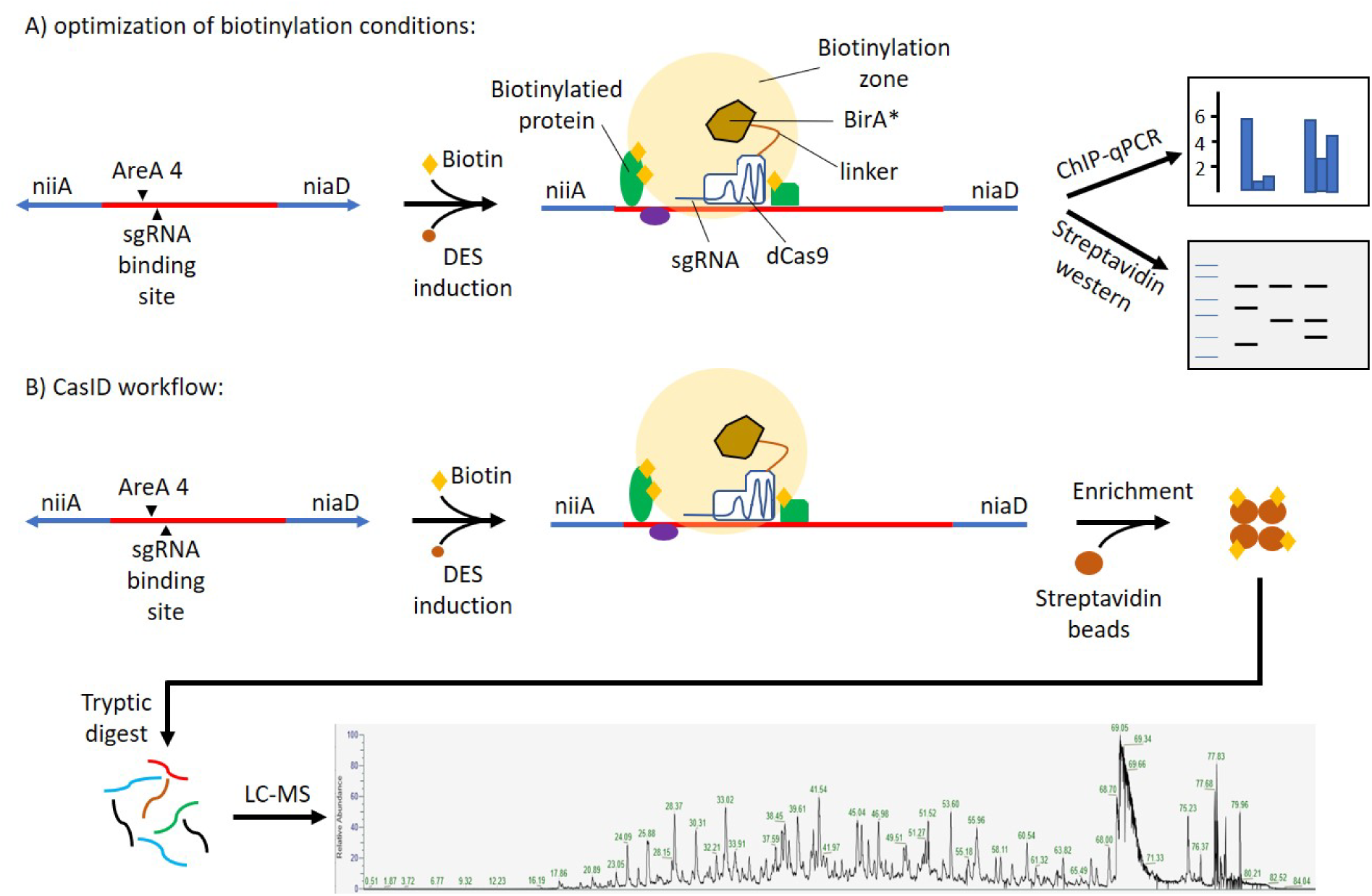
Workflow for optimization of biotinylation conditions (A) and the CasID workflow (B)

### Preparation of the TurboID-dCas plasmids for transformation

As backbone of the TurboID-dCas plasmid, dmWt4-BirA_5 was digested with EcoRI (Schüller et al., 2020). Primers which were used for construction of the plasmid are listed in Supplementary File 1.

#### Construction of sgRNA-nirAII/AreAIV

The sgRNA was constructed analogous to Schüller et al. (2020) using the primers dCas9-gRNA-areA4_nirAII-fwd-Afum and dCas9-gRNA-areA4_nirAII-rev-Afum.

The fragments were annealed using annealing buffer (10 mM Tris, pH 7.5 - 8.0, 50 mM NaCl, 1 mM EDTA) in the PCR cycler. The Plasmids were generated by yeast recombinational cloning and introduced in *A. nidulans* ectopically.

### Proximity biotinylation assay

All tested strains were plated on AMM+NO_3_ and the respective supplements followed by incubation at 37°C for 3 to 5 days. The spores were harvested with 5 ml 0.1% Tween 20 and filtered through glass wool. To homogenize the spores in the suspensions, they were mixed for 6 hours at 4°C on a rotary wheel. In a 1000 mL shake flask 200 mL AMM with appropriate auxotrophic supplements, 10mM proline as neutral N-source for growth, 1nM DES for TurboID-dCas induction were inoculated with 4*10^6^ spores/ mL and incubated at 37°C, 180 rpm for 17 hours. After this initial growth phase, 100 µM biotin was added to provide sufficient biotin for the biotinylation reaction. To modulate AreA activity, NaNO_3_^−^ was added to a final concentration of 150 µM for AreA activation or NH_4_^+^ tartrate to a final concentration of 5 mM for AreA inactivation. Amended cultures were then further incubated for 1h at 37°C, 180 rpm. Finally, the samples were harvested using Miracloth filter (Merck), washed with sterile tap water to remove extracellular biotin and then frozen in liquid nitrogen. The samples were homogenized using Retsch mill (f = 30 s^−1^, 30 sec).

### Protein extraction

For protein extraction 100 mg of the pulverized mycelium were put in a 2 ml tube. Protein extraction buffer: 20 mM HEPES pH 7.5, 5 mM MgCl_2_, 10 mM KCl, 2 mM EDTA, 1 mM DTT (prepare fresh!), 0.1 mM PMSF (in DMSO), 1 µM Leupeptin, 1 µM Pepstatin, 20% Glycerol. 1 ml buffer was added and the cells were lysed using a Bioruptor (3 x 30 seconds, 30 seconds break in between the pulses) followed by centrifugation at 15000 rpm for 10 min at 4°C. The supernatant was transferred to a new tube. To remove previously intracellularly available biotin the buffer was exchanged using Amicon 10 kDa columns according to the manufactureŕs instructions.

### Western blot

Equal amounts of the samples were loaded on a 10% SDS-PAGE and run at 160 V for 1 h. For the transfer of the proteins to the membrane, Trans-Blot® Turbo^TM^ transfer system (BioRad) was used at 25V, 2.5A, 20 min. The membrane was blocked with 3% BSA in TBST overnight followed by washing of the membrane with TBST for 5 minutes. Subsequently the primary antibody (HA, Streptactin-HRP or Cas9) in 3% BSA was added and the samples were incubated for 1h. The membrane was washed three times with TBST for 15 minutes, respectively. The secondary antibody was added and incubated for one hour. The membrane was washed again with TBST three times for 15 minutes, respectively, before developing and visualization.

### Chromatin immunoprecipitation and qPCR

For chromatin immunoprecipitation the CasID test strains, AreA-HA as well as an untagged wild type were used. All tested strains were inoculated in Aspergillus minimal media (AMM) with glucose as C-source, 10 mM proline as N-source as well as the respective supplements. After incubation for 17 hours at 37°C, 180 rpm, 100 µM biotin, 150 µM NaNO_3_ for short-term induction or 10 mM NH_4_ ^+^ for short-term repression as well as 1 nM diethylstilbestrol (DES) for TurboID-dCas induction were added to the medium. These cultures were then incubated for one hour at 37°C, 180 rpm. For crosslinking, formaldehyde (1% final concentration) was added followed by incubation at 20°C, 180 rpm for 15 minutes followed by addition of 125 mM glycine and incubation at 37°C, 180 rpm for 5 minutes. The mycelia were harvested, washed with water to remove excess of biotin and frozen in liquid nitrogen followed by pulverization using Retsch mill MM 400.

For the ChIP ~30 mg of the ground mycelia were put in a tube and 1 ml ChIP lysis buffer (50 mM Hepes-KOH pH 7.5, 140 mM NaCl, 1 mM EDTA pH 8.0, 1% Triton X-100, 0.1% sodium deoxycholate and 0.1% SDS + protease inhibitors) was added. The mycelia were sonicated for 25 minutes (2 minutes pulse, 1 minute break). The samples were centrifuged for one minute followed by transfer of the upper phase to a fresh tube. Protein G agarose (ThermoFisher) was equilibrated with ten-fold volume of ChIP lysis buffer. 40 µl of equilibrated protein G were added to the samples which were incubated at 4°C, 16 rpm, for 1 hour, followed by centrifugation and transfer of the liquid phase to a new tube. The sample was split in 100 µl aliquots to which 900 µl dilution buffer (0.5% Triton X-100, 0.5% sodium deoxycholate, 2 mM EDTA pH 8.0, 20 mM Tris-HCl pH 8.0, 150 mM NaCl) were added. 100 µl of one dilution were taken aside as input control and kept at −80°C until reverse crosslinking. To the remaining 900 µl of the sample 1 µg antibody (Cas9 or HA) was added followed by incubation at 4°C, 16 rpm overnight.

For precipitation of the protein-antibody conjugate, 40 µl Dynabeads^TM^ Protein G for immunoprecipitation (ThermoFisher) were added. The samples were incubated for 40 minutes at 4°C. The tubes were put on a magnetic rack until the beads were settled and the liquid phase was removed. Subsequently, the beads were washed three times with low salt buffer (0.5% Triton X-100, 0.5% sodium deoxycholate, 2 mM EDTA pH 8.0, 20 mM Tris-HCl pH 8.0, 150 mM NaCl) and one time with high salt buffer (0.5% Triton X-100, 0.5% sodium deoxycholate, 2 mM EDTA pH 8.0, 20 mM Tris-HCl pH 8.0, 500 mM NaCl). The beads were resuspended in 125 µl TES buffer (50 mM Tris-HCl pH 8.0, 10 mM EDTA pH 8.0, 1% SDS) and incubated overnight at 65°C, 850 rpm. 120 µl of the ChIP samples and 100 µl of the input control, respectively, were transferred and 8.0 or 9.6 µl TEP buffer (167 mM EDTA pH 8.0, 667 mM Tris-HCl pH 8.0, 150 U/ml proteinase K) were added. After settling of the beads in a magnetic rack the supernatant was transferred to a new tube followed by DNA purification (Quiagen, Mini Elute).

For the qPCR primers ChIP nirA2/areA4 1F (5’-CGCAATGGACGACCGTCATCG-3’) and ChIP nirA2/areA4 1R (5’-ATCAGAACGCTGCCCTGAGC-3’) were used.

### Enrichment of biotinylated proteins and tryptic digest

After extraction, the protein extracts were loaded on Sera-Mag™ SpeedBead Streptavidin-Blocked Magnetic Particles which were previously blocked by washing with 3% BSA in TBST solution. After incubating for one hour on the rotary wheel, unbound proteins were removed followed by different washing steps as described by (Branon et al., 2018): 2x with cold extraction buffer, 1x with cold 1 M KCl, 1x with cold 100 mM Na_2_CO_3_, 1x with 2M Urea in 10 mM Tris pH 8 and 2x with cold extraction buffer without protease inhibitors and PMSF. After washing the beads five times, the beads were suspended in 200 µl buffer containing 100 mM KCl.

The beads were washed three times with 45 µl 100 mM TEAB followed by alkylation and tryptic digest according to the manufactureŕs instructions. Subsequently, the peptides were purified using C18 spin tips. The samples were suspended in 8 µl 0.1% TFA followed by injection of 6 µl to the MS.

## Results

### Construction of the CasID strain and appropriate control strains

To establish the method with the appropriate controls, a CasID test strain which contained TurboID-dCas, the sgRNA and the HA-tagged AreA (CasID +sgRNA) was constructed. As control strains we generated one strain where only AreA was HA-tagged as well as one additional control strain which had the AreA-HA and also the TurboID-dCas construct but which lacked the sgRNA (CasID −sgRNA), that would be necessary to bring TurboID-dCas at the desired locus of the IGR bidirectional nitrate promoter. In addition, a wild type strain lacking all constructs also served as control. This latter strain provides a negative control for all experiments including Westerns, ChIP experiments and proteomics and also gives an indication for natively biotinylated or unspecific proteins that may bind to the affinity column. The HA-AreA strain serves as control for the Western and ChIP analysis to confirm transcription factor presence at the IGR target locus and again for natively biotinylated or unspecific proteins binding to the affinity resin. The CasID −sgRNA control strain covers all aforementioned unspecific proteins, but also reveals proteins that are biotinylated by the TurboID-dCas during all stages from ribosomal generation until nuclear accumulation. In this case we should find an overlap with the wild type and the HA-AreA strain and additionally find novel biotinylated proteins from various compartments of the fungal cell. Finally, the CasID +sgRNA test strain directing the dCas fusion protein to the selected IGR location in vicinity to the AreA-binding site 4 should add the final set of novel proteins that are present and biotinylated specifically at that locus. TurboID-dCas as well as the sgRNA constructs were ectopically integrated. All strains used for the experimental work are listed in Figure 2. They were tested for apparent phenotypes and showed comparable radial growth to the wild type on minimal media containing either nitrate or ammonium (Supplementary Figure 1).

**Figure 2:**
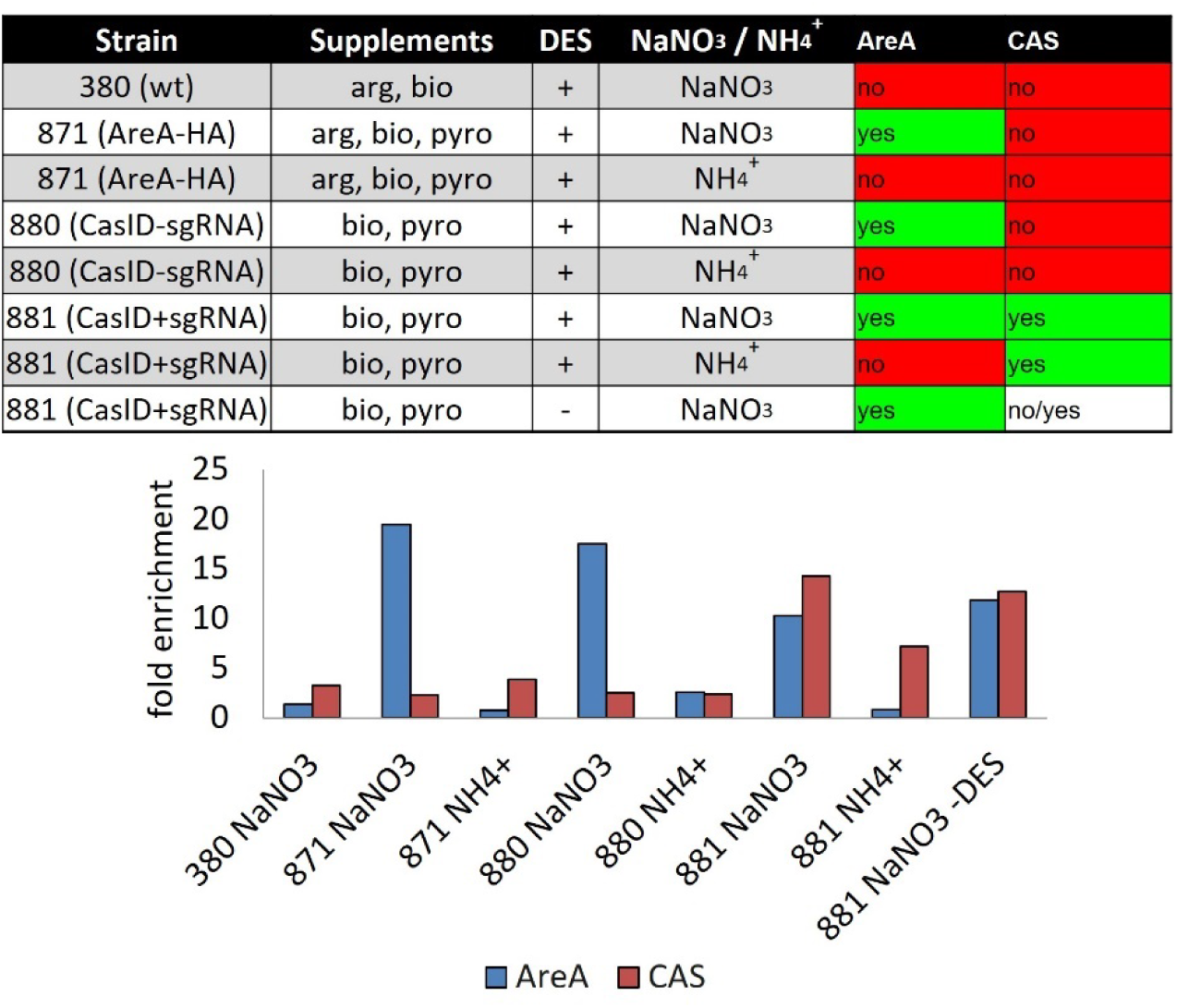
Enrichment of AreA and TurboID-dCas at the nitrate locus under different conditions; 380 wild type (wt) control strain; 871: control strain with HA-tagged AreA; 880: control stain with HA-tagged AreA and CasID-sgRNA; 881: CasID test strain with HA-tagged AreA and CasID+sgRNA; TurboID-dCas and sgRNA directing it to the nitrate promoter 30 bp apart from AreA binding site 4.

A scheme of the workflow and controls for CasID is shown in Figure 1.

### Localization of AreA and CasID under activating and repressing conditions

In order to obtain proof of concept for the method, we wanted to create conditions where AreA as well as our TurboID-dCas fusion protein are both binding to the bidirectional nitrate promoter in close vicinity. This spacial neighborhood was desired as the TurboID part of the fusion protein has a limited functional radius of about 30nm that is mainly provided by the linker region between the TurboID and the dCas peptides (Fig. 1) (May et al., 2020). Therefore biotinylation of AreA by TurboID-dCas would only be feasible when both proteins are positioned next to each other. In our case, a suitable sgRNA PAM-motif is located at a 28 bp - or roughly 10nm - distance from AreA binding site 4 in the IGR (see Fig. 1) and this sequence was therefore chosen as a target locus. Besides the CasID +sgRNA test strain (strain 881) the CasID −sgRNA control strain (strain 880), and the control strain expressing only HA-AreA (strain 871) were used. As negative control for all strains, a wild type strain (strain 380) was grown in parallel. All of these strains were pre-grown on a neutral nitrogen source (proline) and then switched to AreA-activating conditions by addition of 150 µM nitrate and further growth for 1 h. This low nitrate pulse induction for 1 h was chosen to promote AreA binding and activity at the nitrate locus but avoid high levels of the ammonium product that would lead to nitrogen metabolite repression and thus AreA inactivation and export from the nucleus due to the activity of the CrmA exportin (Todd et al., 2005).

Following these consideration, we then performed a set of chromatin immunoprecipitation (ChIP) experiments with all strains to test for the presence of HA-AreA and TurboID-dCas at the nitrate locus. In these ChIPs, AreA was precipitated with an HA-specific antibody and the TurboID-dCas presence was tested using a Cas9-specific antibody. Quantification of binding was carried out by qPCR using a set of primer that encompass both binding sites. **Figure *2*** shows that AreA is strongly enriched at the nitrate locus in the presence of nitrate in all strains that carry the construct (tester strain 881, control strains 880 and 871), but not in the wild type without HA-AreA (strain 380). Whereas AreA is highly enriched in nitrate conditions, it is much less abundant when ammonium served as nitrogen source. This confirms our previous data and also proves that the experimental set-up is correct for the subsequent CasID development. ChIP results for TurboID-dCas binding at the nitrate promoter were only positive in the CasID test strain (881) that contains the TurboID-dCas plus the sgRNA, but not in the control strain (880) where no sgRNA is expressed. As expected, the TurboID-dCas protein was present in the 881 test strain at the nitrate locus under both nitrate-inducing and ammonium-repressing conditions as the TurboID-dCas binding is not expected to be nitrogen source dependent. Interestingly, the estrogen-dependent transcription of the TurboID-dCas gene seems to be leaky in the actual integration locus as binding of the fusion protein was detected in comparable amounts regardless if DES was added to the culture or not (compare lanes 881 NaNO_3_ and 881 NaNO_3_-DES in Fig. 3 (left)). In summary, these ChIP experiments provided strong evidence that the system works as expected in the CasID test strain and all control strains provide appropriate experimental controls to move forward with the method development.

**Figure 3:**
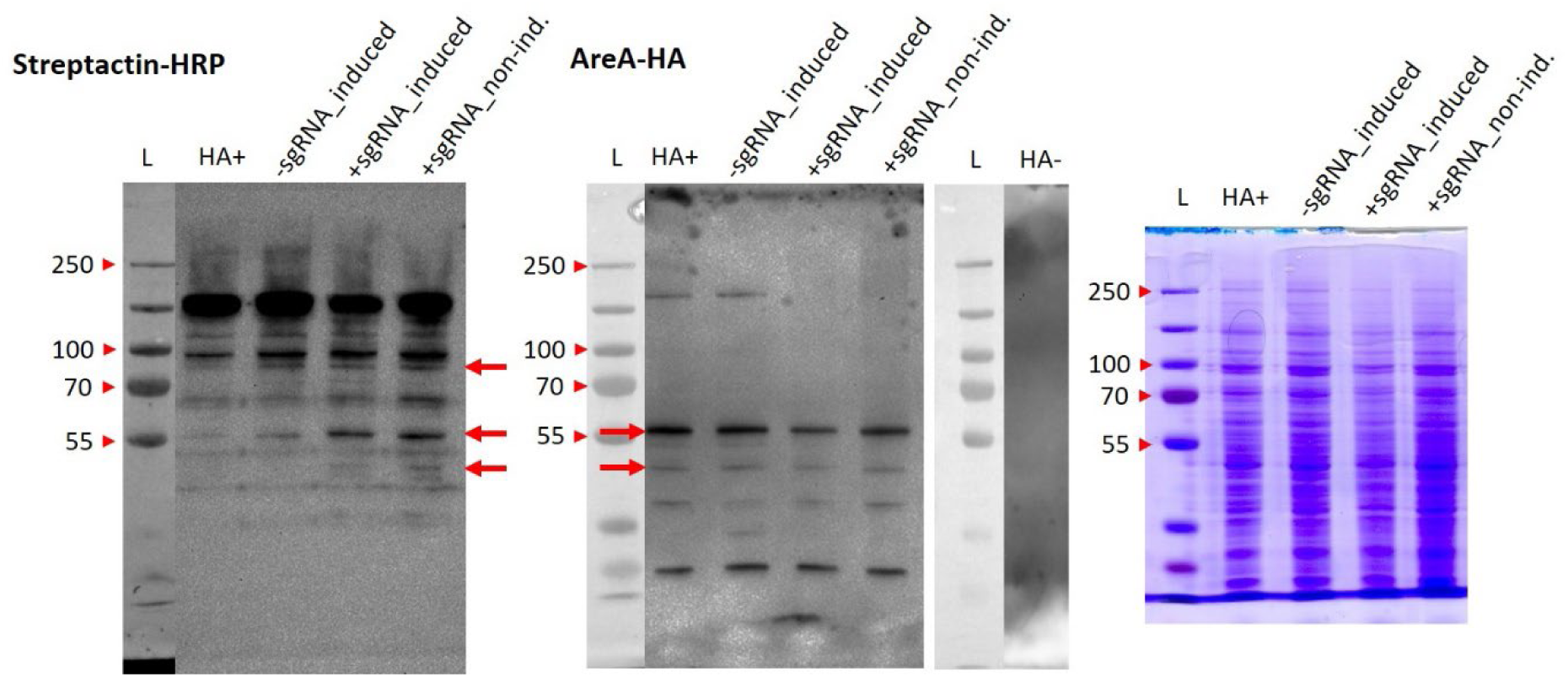
Western blots for the detection of biotinylated proteins via streptactin-HRP (left), the HA.tagged AreA (middle) and a coomassie stained SDS-PAGE as loading control (right). HA+: strain 871 containing only the HA-AreA construct; −sgRNA_induced: strain 880 containing TurboID-dCas without sgRNA grown in the presence of 10nM DES inducer for the TurboID-dCas construct; +sgRNA_induced: strain 881 containing TurboID-dCas and also the sgRNA targeting the construct to the nitrate locus, grown in the presence of 10nM DES; +sgRNA_non-ind.: strain 881 containing TurboID-dCas and also the sgRNA, grown without inducer for the TurboID-dCas construct (note that expression of the construct is leaky and also occurs to some extent in the absence of inducer)

### Optimization of biotinylation conditions and Western analysis of products

With the confirmation that both proteins are at the locus of interest close to each other one hour after induction, biotinylation of AreA may occur. Since all strains except the wild type control contain an HA-tagged AreA, we were able detect this protein by western blots and at the same time probe for biotinylation using a streptactin-HRP detection system. We reasoned that AreA may be biotinylated in the CasID samples and hence there might be an overlap between the HA-derived signal for AreA and a band from the biotinylation assay. In the western blot in **Figure *3***, left panel (streptactin-HRP) biotinylated proteins are revealed. There are several bands visible in all strains (e.g. strong band at around 130 kDa) indicating that these proteins are natively biotinylated in *A.nidulans*. But there are also a few bands visible exclusively in the samples derived from TurboID-dCas containing strains (880, 881). This indicates that the TurboID-dCas fusion protein is functional and biotinylates closely spaced proteins via the Turbo-BirA enzyme. Most of these additional bands appear in the biotinylation assay regardless whether the fusion protein is targeted to the locus (CasID strain 881) or not (control strain 880). However, there are two bands at an apparent size between 30 kDa and 55 kDa visible in this Western blot that specifically appear only in the CasID strain with the sgRNA indicating a successful biotinylation reaction at the nitrate locus. None of these bands, however, would correspond to the expected full size of an AreA monomer of 94 kDa.

The HA-specific Western for the detection of the corresponding AreA band, however, did not show a signal at the expected size of 94 kDa but four signals with lower molecular weight starting at 55 kDa and lower appeared in the HA-AreA carrying strains. These signals were not present in the wild type control (lane HA- in Fig 3, central panel) and thus appear to be specific processing or degradation products of HA-AreA. Interestingly, two of the four bands (55 kDa and slightly below at around 50kDa) are also strongly stained on the streptactin western blot in the CasID strain (labelled +sgRNA) and weakly in the control strain with TurboID-dCas without sgRNA (labelled −sgRNA). Although these similarities between the two western blots could indicate biotinylation of HA-AreA (before processing or degradation), similar sizes of the signals could also be pure coincidence and thus it is advisable to avoid any conclusions before an independent method, e.g. mass spectrometry analysis of biotinylated proteins, would have confirmed these data.

### Enrichment and protein mass spectrometry analysis of biotinylation of products

To obtain a comprehensive view on specific proteins present at the chromatin of the selected locus, the method of choice is protein mass spectrometry of protein extracts that were enriched for biotinylated proteins by affinity purification. The detailed steps of the procedure are described in the Materials and Methods section, but briefly, proteins were extracted from mycelia of the CasID strain and from the control strains using a standard native protein extraction protocol. After this step, protein extracts were loaded on streptavidin coated magnetic beads and washed extensively using different buffers to elute non-specifically attaching proteins from the streptavidin-coated matrix. Optimally, after these washing steps, only biotinylated proteins should remain bound to the beads. Because of the exceptionally high affinity between biotin and streptavidin (K_d_ ~ 10^−14^ M) elution of the bound proteins is only feasible under chemically denaturing conditions and high temperatures (Deng et al., 2013). Moreover, it is known that biotinylation of lysine drastically reduces the efficiency of trypsin to cleave after the biotinylated lysine residue (Li et al., 2021). To circumvent these problems, a two-step elution of streptavidin-bound proteins was pursued. In the first step, beads were subjected to tryptic digest directly in the suspension to release peptides that can be cut at arginines or non-biotinylated lysines. In the second step, the remaining peptides bound via their biotinylated lysine(s) to the streptavidin-coated beads were eluted using excess of biotin and denaturing, but still MS-compatible conditions. Both elution samples were subsequently combined and subjected to MS analysis.

The LC-MS analysis was done from two biologically and technically independent extractions and enrichments. To our surprise, the number of total proteins detected in all control strains, was quite high and was in the range between 800 – 900 different proteins detected already in the control samples. Only in the CasID strain, the number of total proteins detected by MS was slightly higher and in the range between 1.100 – 1.400 different proteins (Table 1). These MS results indicated that despite several rigorous washing steps, the number of proteins unspecifically attaching to the streptavidin-coated beads is very high. Several attempts using different buffers and washing conditions based on published protocols and a literature search revealed that this problem existed also in many other studies using this enrichment method (reviewed in (Trinkle-Mulcahy, 2019). Therefore, a set of rigorous genetic and experimental controls is necessary that allows to discriminate the specific proteins labelled by the TurboID-dCas enzyme from the general background during enrichment.

**Table 1:** Summary of identified proteins with and without biotinylation; AreA-HA is the HA-tagged control strain, CasID-sgRNA contains the biotin ligase fused to cas9 without sgRNA (background biotinylation), CasID+sgRNA also contains a sgRNA.

| Sample name | Number of identified proteins w/o biotinylation | Number of identified proteins with biotinylation | Total Number of identified proteins |
| --- | --- | --- | --- |
| AreA-HA_1 | 916 | 3 | 919 |
| CasID-sgRNA_1 | 983 | 5 | 988 |
| CasID+sgRNA_1 | 1119 | 5 | 1124 |
| AreA-HA_2 | 909 | 7 | 916 |
| CasID-sgRNA_2 | 852 | 2 | 854 |
| CasID+sgRNA_2 | 1433 | 4 | 1437 |

In the two independent measurements of each strain a partially different protein profile was detected, however, the majority of the proteins within the replicates was identical in both biologically independent experiments.

Surprisingly, proteins carrying a biotinylation were hardly detected in any sample. **Figure 4A** shows a Venn diagram derived from Table 1 where the proteins carrying a biotin moiety from the different strains are visualized. We found six “natively biotinylated” proteins that appeared in all analyzed strains at least in one of the two replicates (**Figure 4B**). For some of these proteins, like biotin carboxylase or pyruvate carboxylase, biotinylation is a known posttranslational modification. Others, like AN4308 (predicted translation protein), AN3636 (predicted phosphatidyl-inositol phospholipase C, an enzyme known to be involved in signal transduction) or AN1394 (predicted septin type G-protein cytoskeletal protein) are so far not known to carry biotin as a PTM and the significance of this finding needs to be validated by additional repetitions testing also other genetic backgrounds.

**Figure 4:**
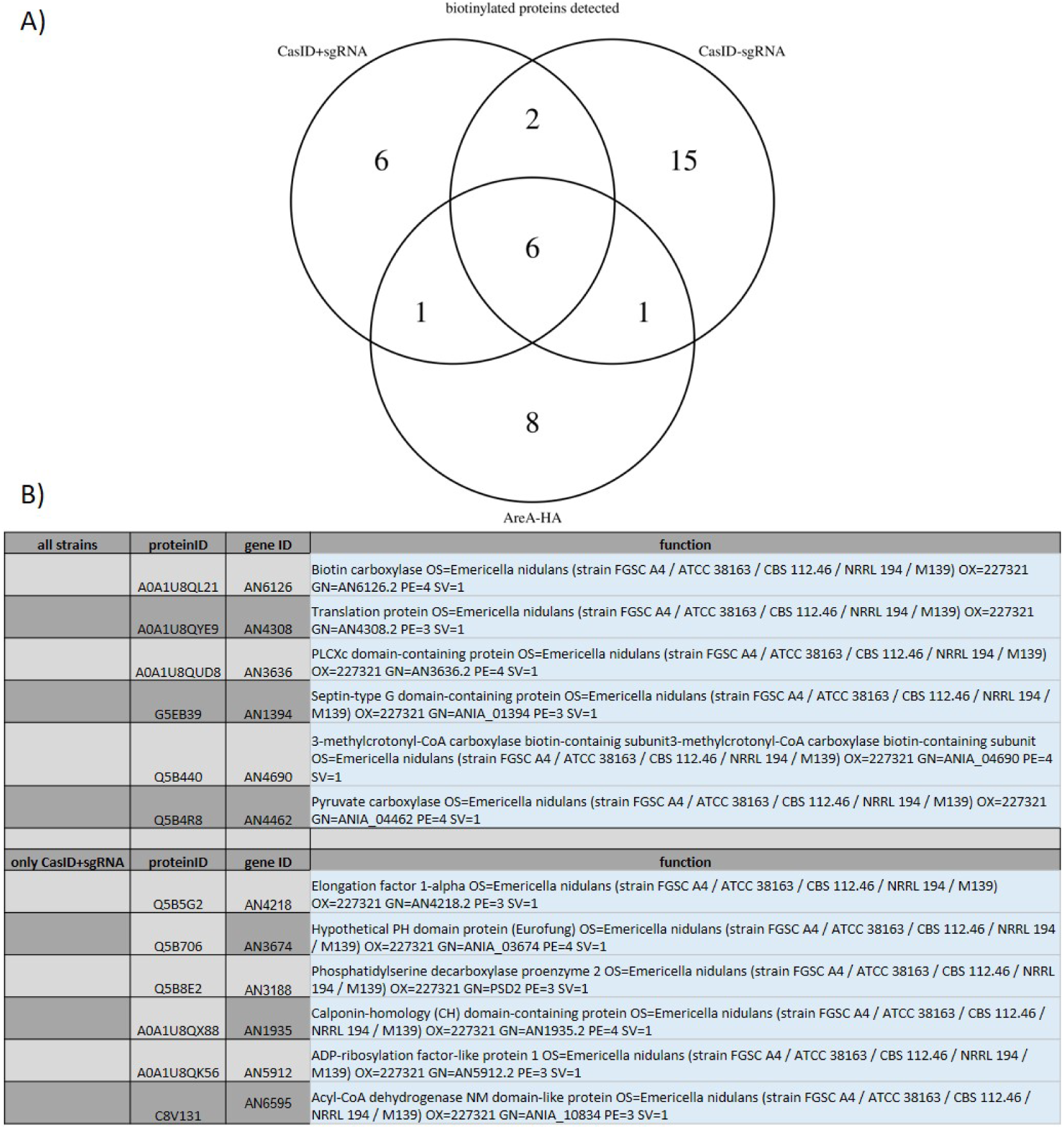
Venn diagrams of biotinylated proteins identified in at least one of the two independent measurements of the strains

Another six proteins appeared to be biotinylated exclusively in the CasID test strain expressing TurboID+sgRNA. In this population of biotinylated peptides we expected mainly proteins that are known to associate with chromatin. The identity of these proteins, however, is quite divers.

For example, we find biotinylated elongation factor 1 alpha (gene AN4218) that could have been biotinylated during the process of translating TurboID-dCas9. A calponin-homology domain containing protein (gene AN1935), known for its actin binding capacities (Yin et al., 2020), could have been at the chromatin locus, as well as AN3674, a pleckstrin-homology domain (PH) protein that is also related to cytoskeletal function and membrane targeting and AN5912, that encodes a predicted GTPase regulating vesicle formation during intracellular traffic and also related to processes in which PH proteins are involved. (Lemmon, 2004) On the other hand, there is no obvious “chromatin-connection” to the AN3188 product that is predicted to be involved in phospholipid metabolism, and AN6595 that codes for a predicted acylCoA-DH enzyme involved in peroxisomal beta oxidation of fatty acids. Overall, these six proteins for which biotinylated peptides have been identified by MS at least in one of the experiments have some known chromatin-related functions but each of these proteins should be independently validated by tagging, purification and MS analysis identifying the biotinylation status. rather represent proteins that were biotinylated during the transit of the TurboID-dCas9 protein from the site of translation to the nucleus and further to the target site at the nitrate promoter.

In order to improve the selectivity of our analyses for specific proteins only appearing in the CasID+sgRNA strain we restricted all MS peptide hits to those that appeared in both independent measurements of each strain, but regardless if these proteins are biotinylated or not as the biotin moiety may still be bound to the streptavidin beats after the tryptic digest and also not been eluted in the second step (Figure 5).

**Figure 5:**
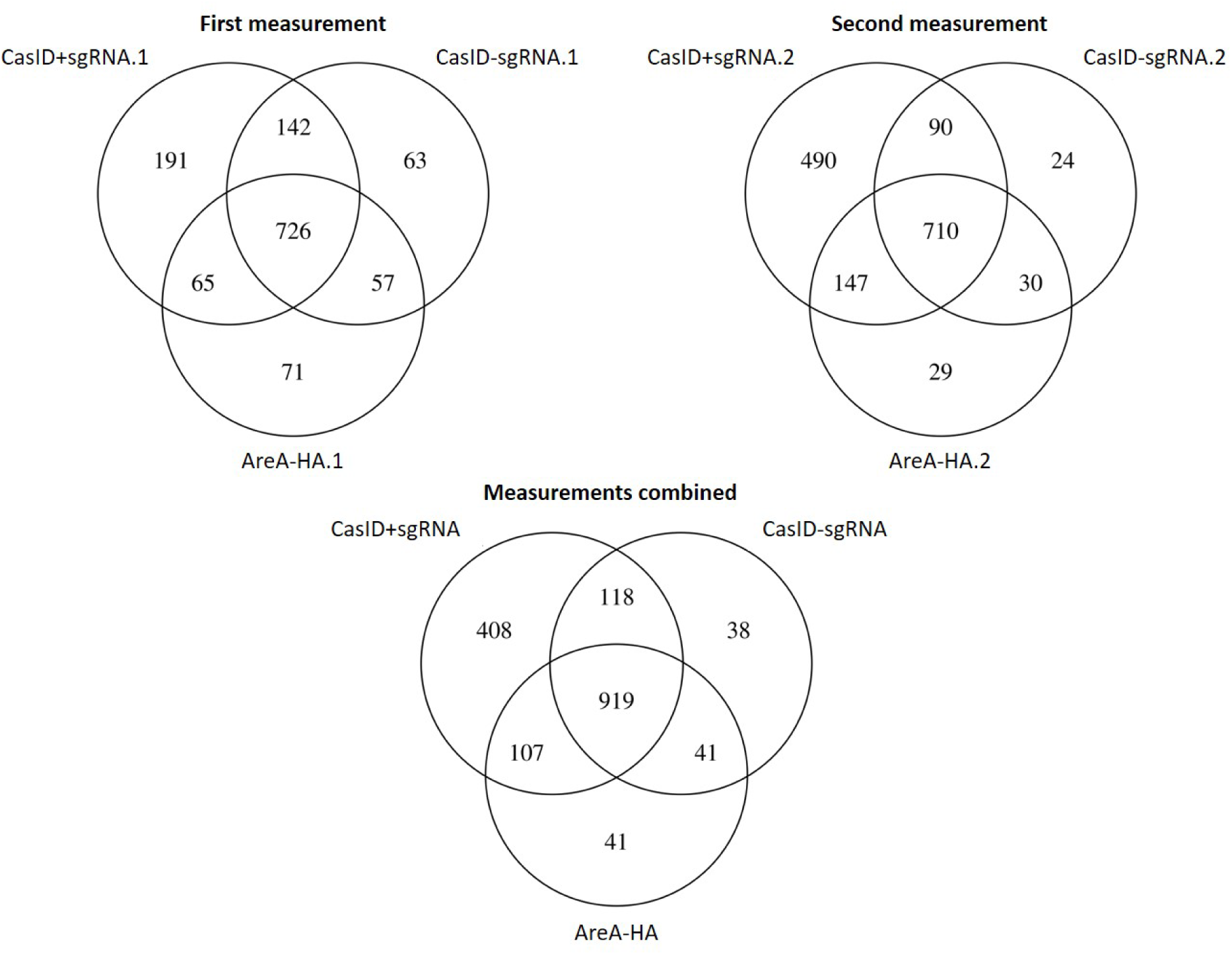
Venn diagrams of proteins identified in the two independent measurements of the strains and of both measurements combined

While in the first measurement only 191 proteins were exclusively present in the CasID+sgRNA strain, in the second approach 490 proteins were identified. After merging the protein IDs of both measurements 408 proteins appeared at least once and only 32 of them were detected in both measurements specifically in the CasID+sgRNA strain. A complete list of these 32 consistently detected proteins with their proposed function is shown in Table 2. There are several putative and uncharacterized proteins included but also a few with an already described function. The accessions highlighted in red font are known to be located in the nucleus. Among the already characterized proteins are PhnA (gene ANIA_00082), a phosducin like protein required for vegetative growth, developmental control, and toxin biosynthesis, BudA (gene ANIA_01324) involved in septum formation, CmkB (gene ANIA_03065) a calmodulin dependent protein kinase. Strikingly, we also found NmrA (gene ANIA_08168), a regulatory protein negatively regulating AreA activity. NmrA interacts with AreA at its C- as well N-terminus and thereby blocks its activity und nitrogen metabolite repressive conditions. AreA itself was not among the proteins falling into the selection category (only in the CasID+sgRNA in both independent repetitions), although it was detected in one of the experiments in this strain.

**Table 2:**
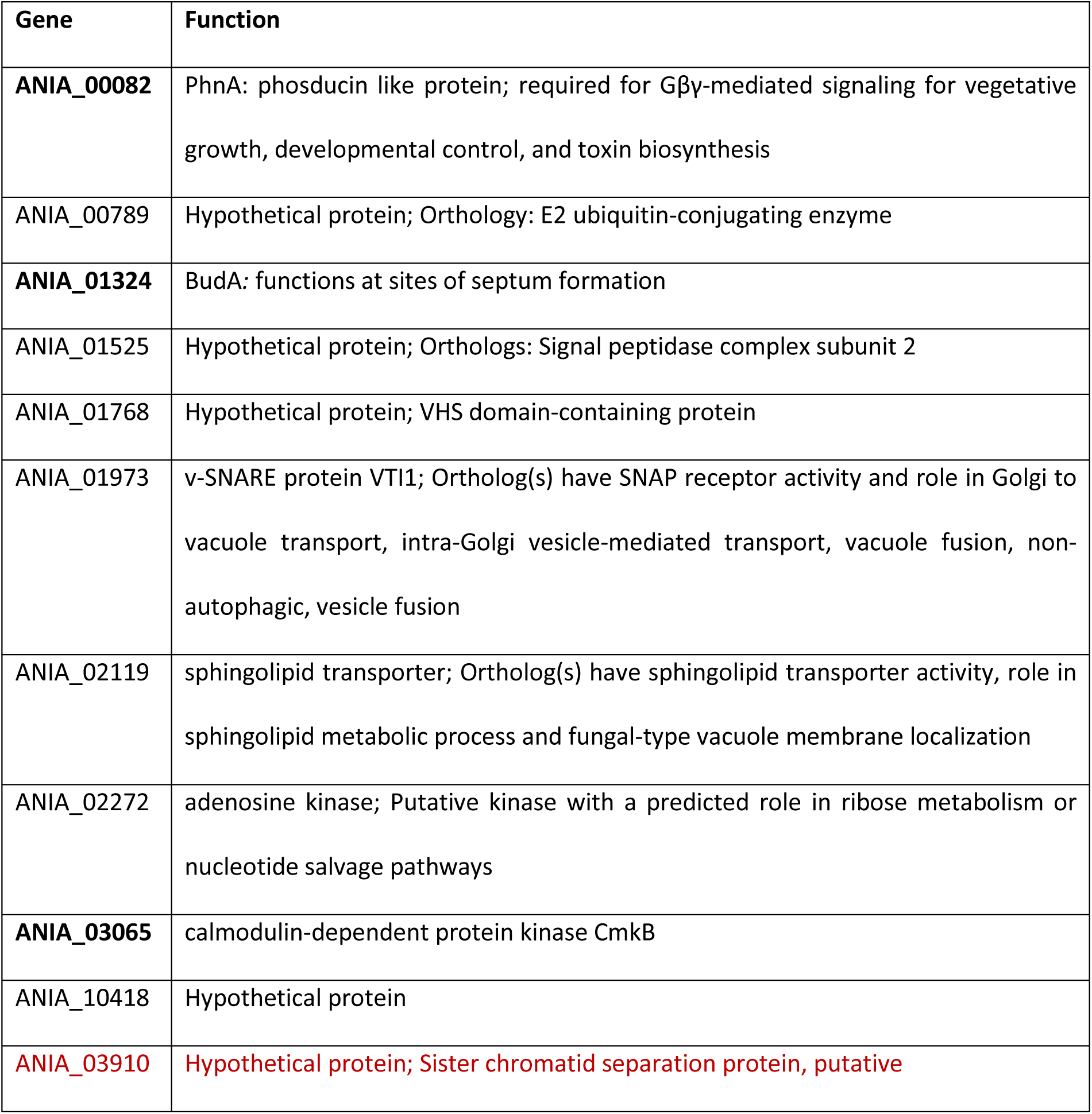
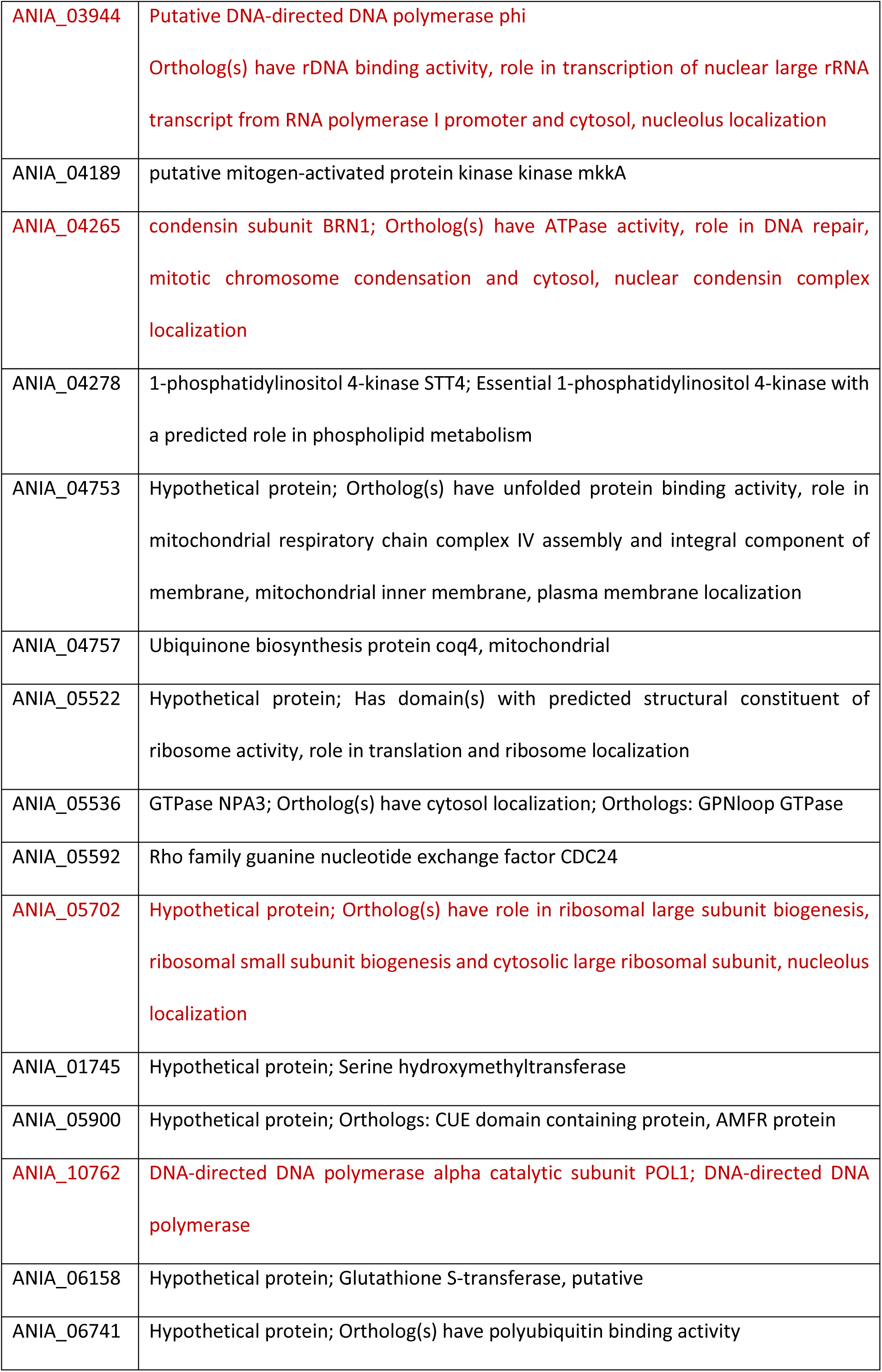
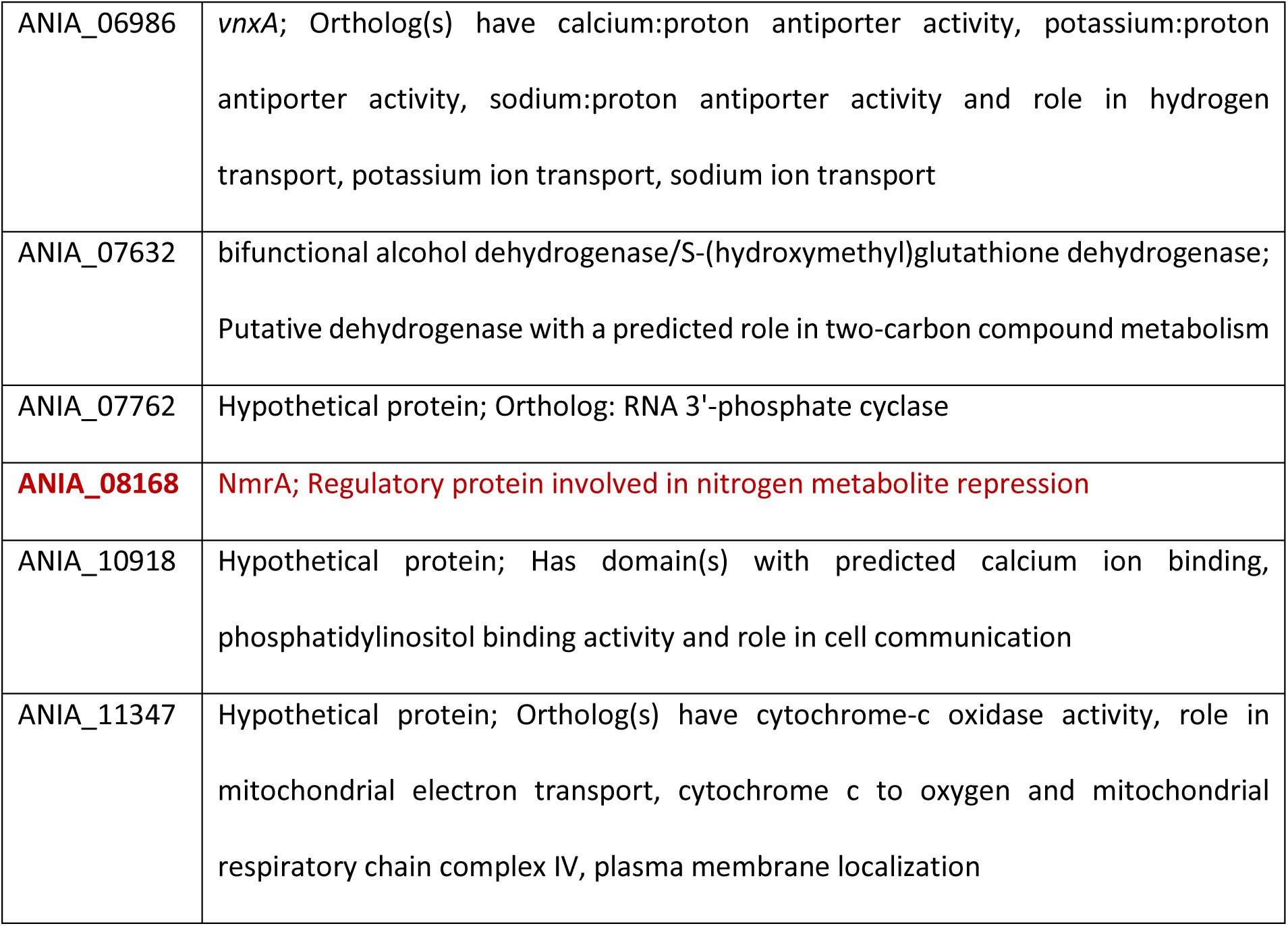
Genes where corresponding proteins were exclusively identified in the CasID strain; bold = characterized proteins; red = known nuclear proteins.

## Discussion and Conclusions

For method development we targeted the fusion enzyme to the well characterized nitrate locus. This locus was chosen due to the possibility of rigorous controls and the possibility to test CasID and an adjacent regulator binding first by ChIP to see if the system works. MS results of the locus CasID confirmed the ChIP data in two ways: (i) we found several chromatin-related proteins specifically only in the CasID+sgRNA strain indicating that biotinylation occurred at the locus, and (ii) although we did not consistently find our main target AreA as biotinylated protein in MS analysis, we found the AreA-interacting regulator NmrA exclusivey in the CasID+sgRNA strain. At this point we can only speculate on the significance of this finding, but it could be that the TurboID part of the fusion protein could reach out and target the interacting NmrA, but not the “underlying” AreA regulator itself. In any case, the result fits well with the known function of AreA regulation by NmrA. In addition, our data would indicate that NmrA modulates AreA activity while always interacting with it, as we found this interaction during nitrate conditions. This would mean that NmrA was already present at the locus although not negatively interfering with AreA function. Future studies may be warranted that look into this interaction in more detail. It could be that the biotinylation radius with the biotinyl-5’-AMP cloud around the TurboID of CasID was too narrow to find also AreA. One possibility to enlarge the biotinylation radius would be the introduction of a longer linker between dCas9 and TurboID allowing more flexibility. Since DNA and chromatin are flexible structures, not static, it is also possible that AreA was simply out of reach or that NmrA was closer to CasID and preventing AreA from being biotinylated. In addition to a longer linker between sCas9 and TurboID, also the employment of more than one sgRNAs per locus would be an alternative (“primer walking”). In our case, in the bidirectional promoter between *niiA* and *niaD* there are ten binding sites for AreA and four binding sites for NirA described (Muro-Pastor et al., 1999). By using this“primer walking” strategy setting sgRNAS around all AreA binding sites more chromatin regulators of this locus could be identified.

Other characterized proteins consistently detected only in the CasID+sgRNA test strain at the selected nitrate locus were were PhnA, BudA, or CmkB. PhnA is a phosducin-like protein which are described as regulators of G-protein signaling disrupting the G-protein interaction of subunit Gα and Gβγ (Blüml et al., 1997), and the predicted localization of PhnA using “WoLF PSORT Prediction” (https://www.genscript.com/wolf-psort.html) is with the highest probability in the nucleus. Deletion of the gene in *A. nidulans* results in reduced biomass, asexual sporulation in liquid submerged culture, and defective fruiting body formation. Also mycotoxin production is strongly affected since *aflR* is no longer transcribed in the deletion strain (Seo and Yu, 2006). Since we also detected PhnA here by CasID at the nitrate locus it may play a general role in chromatin function. BudA belongs to the actin-binding proteins and has been found in *A.nidulans* to function at sites of septum formation (Harris et al., 2009; Virag and Harris, 2006). So presumably, this protein is not expected to reside at the nitrate locus, however, fungi feature a special type of “open mitosis” in which the cytoskeleton, including actin, reaches through the partially disassembled nuclear envelope to establish intimate contacts with the segregating chromosomes (Boettcher and Barral, 2013). Mutants of Bud6, the BudA homolog of *S.cerevisiae*, show defects in mitosis, possess fewer secretory vesicles, form abnormal septa, and display abnormal actin bars in nuclei. As we also found cytoskeleton-associated proteins in the list of biotinylated proteins specific for the CasID+sgRNA strain, we still need to consider BudA as true chromatin-interacting protein captured at our target locus most likely during mitosis. Also the third characterized protein identified in this CasID screen, CmkB has a role in mitosis and nuclear division. It is a calmodulin-dependent kinase needed for the proper temporal activation of the main cell cycle kinase NimX (CDC2) and timely progression of the nuclear division (Joseph and Means, 2000).

Overall, it seems that the our CasID strategy was successful in capturing an expected regulator at the nitarte locus (NmrA) and several other chromatin-associated proteins such as a condensin subunit (ANIA 04265), DNA polymerase (ANIA 03944), sister chromatid separation (ANIA 03910) and other proteins that regulate mitosis and cell division. This high proportion of cell-cycle-associated proteins might be due to the fact that our biotin-labelling time was over a period of roughly 12 hours and thus several mitoses have certainly occurred during this time. This could lead to the observed biotinylation of cytoskeletal proteins, as moreover, they are highly abundant in the nucleus during the cell division process. As these proteins are most likely not really specific for the target locus, it will be necessary in the future CasID approaches to shorten the labelling time of CasID at the locus. As we have a conditional expression system for the TurboID-dCas9 construct and can also add the additional biotin for a shorter time period, such time restriction of the biotinylation process is feasible. This adaptation may lead to a more locus-specific labelling of the chromatin-associated proteome. Another abundant family of detected prtoeins are related to membrane biogenetics and trafficking. This is also not suprising, as it is known that mitotic exit (Davies et al., 2004) or active transcription locates large genomic regions in close vicinity to the nuclear membrane and thus establishing an intimate connection between membrane biology and the 3-dimensional genome architecture (Sosa Ponce et al., 2024).

Certainly, a strong bottleneck of the CasID method is the enrichment of biotinylated proteins via the strepatavidin-cated paramagnetic beads. This procedure leads to a high level of unspecific proteins that adhere to the matrix without biotinylation and it has already been reported many times in the literature, that some of these “contaminants” can not be eliminatedeven by extensive washing with compatible buffer systems. Ideally, we would get a few biotinylated proteins, however, in every measurement and sample beyond 800 proteins could be unambiguously identified. The high background after enrichment has previously been documented as a challenge in several studies (Branon et al., 2018; Mair et al., 2019; Zhang et al., 2020). As shown in Supplementary Figure 2 the majority of the proteins were detected in both of the two independent measurements in the respective strains. For example, 727 proteins were detected in both measurements in the AreA-HA control strain which does not have the TurboID-dCas9 biotin ligase. In comparison, there were 1009 proteins identified in both measurements of our CasID+sgRNA test strain. After subtracting proteins which were identified in all three control strains, only 32 proteins remained exclusively and consistently detected in the CasID test strain. These results point out once more how crucial appropriate controls are in proximity labelling experiments.

The goal of this study was to adapt the CasID method to identify chromatin-related proteomes at a specific locus of interest in our model organism *A. nidulans*. One type of the loci to be studied in future will be biosynthetic gene clusters (BGCs) encoding secondary metabolites. These genomic regions seem to have a very specific chromatin landscape that is far from being understood at the molecular level (Gacek and Strauss, 2012; Pfannenstiel and Keller, 2019) and thus CasID could help in identifying new regulators. In all fungi studied so far, the majority of genomic clusters coding for secondary metabolites (biosynthetic gene clusters, BGCs) are usually silenced by default during active growth through heterochromatin structures (Atanasoff-Kardjalieff and Studt, 2022; Connolly et al., 2013; Reyes-Dominguez et al., 2012; Soyer et al., 2014; Studt et al., 2016) and are only activated in response to environmental or developmental signals but then often lack typical “active” chromatin signatures like histone H3-lysine 4 methylation (Gacek-Matthews et al., 2016; Karahoda et al., 2022). Thus far, BGC-specific components at a certain locus can only be studied by ChIP provided specific antibodies are available for known posttranslational modifications or tagged proteins. CasID may help to identify new regulatory components at loci of interest as it can be employed to identify parts of the condition-specific proteome in a high spatial resolution using different sgRNAs for targeting the dCas-BirA fusion enzyme. Nevertheless, we need to be aware that this method can give us a first information on proteins potentially present at a locus and additional verification and confirmation is indispensable.

## Acknowledgments

This work was supported by grant Nr. P32790-B of the Austrian Science Fund FWF to JS.

## Author contributions

TS – experimental design and execution, data evaluation, preparation of the draft manuscript; DNL – experimental design and execution, data evaluation; AS – Preparation of the test strains; KH, SS and ERF – proteomics analysis; JS – Experimental design, data analysis and evaluation, funding acquisition

---

**Supplementary Figure 1:**
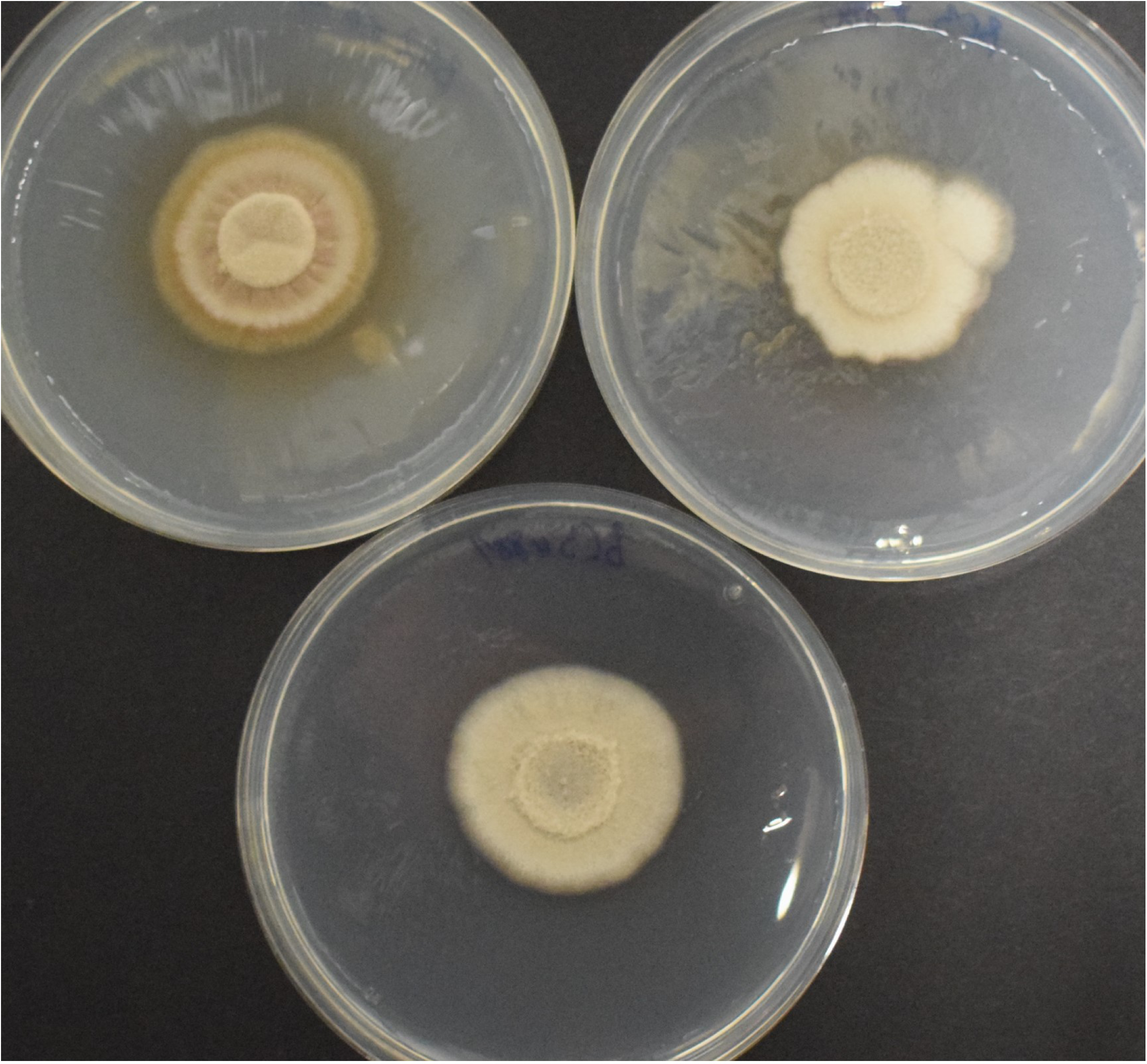
Phenotypic appearance of the tested strains; AreA-HA 3slD-sgRNA (top, right), CaslD + sgRNA (bottom)

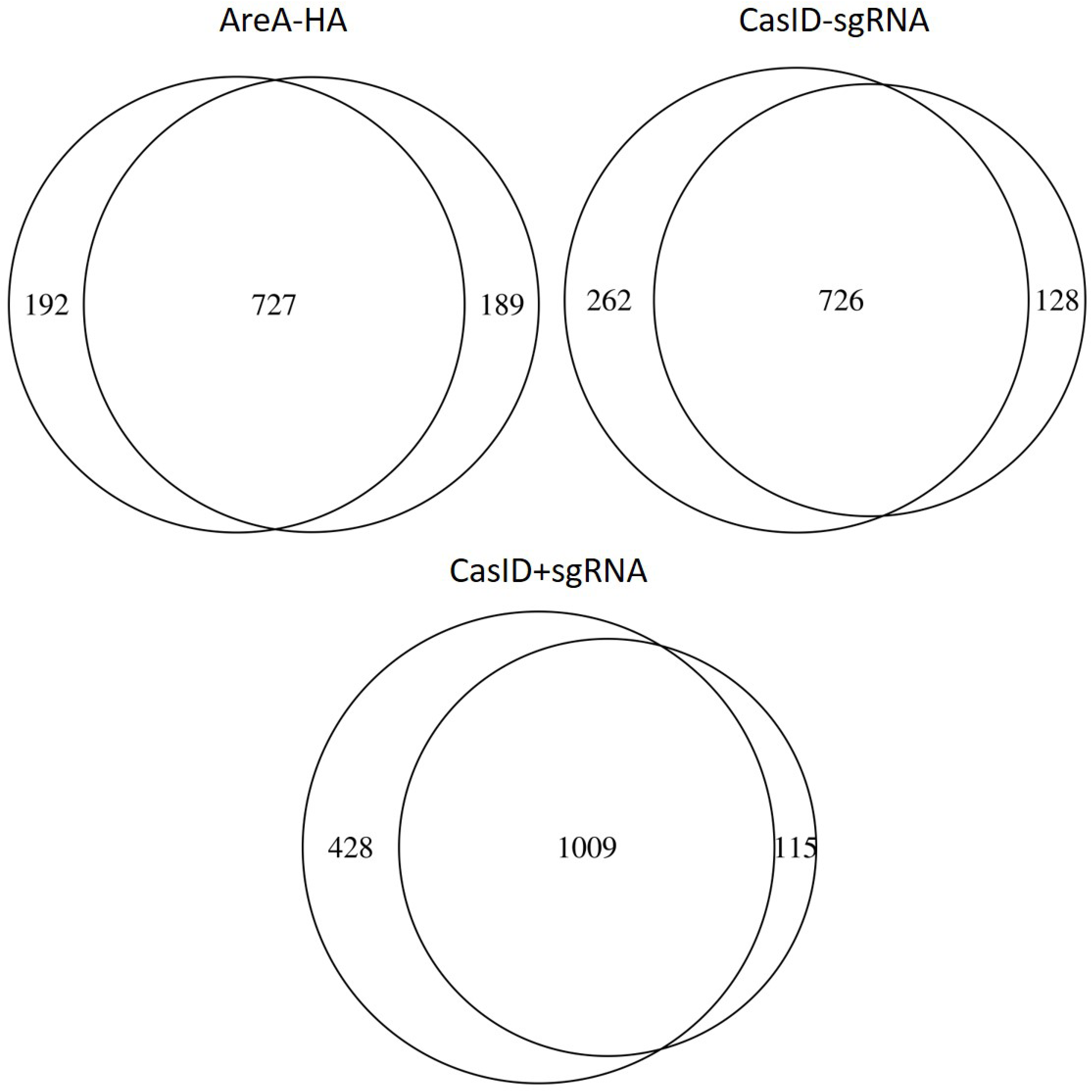

## References

Anton, T., Karg, E., Bultmann, S., 2018. Applications of the CRISPR/Cas system beyond gene editing. Biol. Methods Protoc. 3, bpy002. 10.1093/biomethods/bpy002

Blüml, K., Schnepp, W., Schröder, S., Beyermann, M., Macias, M., Oschkinat, H., Lohse, M.J., 1997. A small region in phosducin inhibits G-protein betagamma-subunit function. EMBO J. 16, 4908– 4915. 10.1093/emboj/16.16.4908

Branon, T.C., Bosch, J.A., Sanchez, A.D., Udeshi, N.D., Svinkina, T., Carr, S.A., Feldman, J.L., Perrimon, N., Ting, A.Y., 2018. Efficient proximity labeling in living cells and organisms with TurboID. Nat. Biotechnol. 36, 880–887. 10.1038/nbt.4201

Caesar, L.K., Kelleher, N.L., Keller, N.P., 2020. In the fungus where it happens: history and future propelling Aspergillus nidulans as the archetype of natural products research. Fungal Genet. Biol. FG B 144, 103477. 10.1016/j.fgb.2020.103477

Cho, K.F., Branon, T.C., Udeshi, N.D., Myers, S.A., Carr, S.A., Ting, A.Y., 2020. Proximity labeling in mammalian cells with TurboID and split-TurboID. Nat. Protoc. 15, 3971–3999. 10.1038/s41596-020-0399-0

Drott, M.T., Bastos, R.W., Rokas, A., Ries, L.N.A., Gabaldón, T., Goldman, G.H., Keller, N.P., Greco, C., 2020. Diversity of Secondary Metabolism in Aspergillus nidulans Clinical Isolates. mSphere 5, e00156–20. 10.1128/mSphere.00156-20

Gawlik, J., Koper, M., Bogdanowicz, A., Weglenski, P., Dzikowska, A., 2022. Nuclear Functions of KaeA, a Subunit of the KEOPS Complex in Aspergillus nidulans. Int. J. Mol. Sci. 23, 11138. 10.3390/ijms231911138

Han, X., Qiu, M., Wang, B., Yin, W.-B., Nie, X., Qin, Q., Ren, S., Yang, K., Zhang, F., Zhuang, Z., Wang, S., 2016. Functional Analysis of the Nitrogen Metabolite Repression Regulator Gene nmrA in Aspergillus flavus. Front. Microbiol. 7, 1794. 10.3389/fmicb.2016.01794

Inglis, D.O., Binkley, J., Skrzypek, M.S., Arnaud, M.B., Cerqueira, G.C., Shah, P., Wymore, F., Wortman, J.R., Sherlock, G., 2013. Comprehensive annotation of secondary metabolite biosynthetic genes and gene clusters of Aspergillus nidulans, A. fumigatus, A. niger and A. oryzae. BMC Microbiol. 13, 91. 10.1186/1471-2180-13-91

Kong, Q., Chang, P.-K., Li, C., Hu, Z., Zheng, M., Sun, Q., Shan, S., 2020. Identification of AflR Binding Sites in the Genome of Aspergillus flavus by ChIP-Seq. J. Fungi Basel Switz. 6, 52. 10.3390/jof6020052

Lamb, H.K., Leslie, K., Dodds, A.L., Nutley, M., Cooper, A., Johnson, C., Thompson, P., Stammers, D.K., Hawkins, A.R., 2003. The negative transcriptional regulator NmrA discriminates between oxidized and reduced dinucleotides. J. Biol. Chem. 278, 32107–32114. 10.1074/jbc.M304104200

Lamb, H.K., Ren, J., Park, A., Johnson, C., Leslie, K., Cocklin, S., Thompson, P., Mee, C., Cooper, A., Stammers, D.K., Hawkins, A.R., 2004. Modulation of the ligand binding properties of the transcription repressor NmrA by GATA-containing DNA and site-directed mutagenesis. Protein Sci. Publ. Protein Soc. 13, 3127–3138. 10.1110/ps.04958904

Mair, A., Xu, S.-L., Branon, T.C., Ting, A.Y., Bergmann, D.C., n.d. Proximity labeling of protein complexes and cell-type-specific organellar proteomes in Arabidopsis enabled by TurboID. eLife 8, e47864. 10.7554/eLife.47864

Muro-Pastor, Mi., Gonzalez, R., Strauss, J., Narendja, F., Scazzocchio, C., 1999. The GATA factor AreA is essential for chromatin remodelling in a eukaryotic bidirectional promoter. EMBO J. 18, 1584–1597. 10.1093/emboj/18.6.1584

Pan, H., Feng, B., Marzluf, G.A., 1997. Two distinct protein-protein interactions between the NIT2 and NMR regulatory proteins are required to establish nitrogen metabolite repression in Neurospora crassa. Mol. Microbiol. 26, 721–729. 10.1046/j.1365-2958.1997.6041979.x

Pateman, J.A., Cove, D.J., 1967. Regulation of nitrate reduction in Aspergillus nidulans. Nature 215, 1234–1237. 10.1038/2151234a0

Punt, P.J., Strauss, J., Smit, R., Kinghorn, J.R., van den Hondel, C.A., Scazzocchio, C., 1995. The intergenic region between the divergently transcribed niiA and niaD genes of Aspergillus nidulans contains multiple NirA binding sites which act bidirectionally. Mol. Cell. Biol. 15, 5688–5699. 10.1128/MCB.15.10.5688

Samavarchi-Tehrani, P., Samson, R., Gingras, A.-C., 2020. Proximity Dependent Biotinylation: Key Enzymes and Adaptation to Proteomics Approaches *. Mol. Cell. Proteomics 19, 757–773. 10.1074/mcp.R120.001941

Schüller, A., Wolansky, L., Berger, H., Studt, L., Gacek-Matthews, A., Sulyok, M., Strauss, J., 2020. A novel fungal gene regulation system based on inducible VPR-dCas9 and nucleosome map-guided sgRNA positioning. Appl. Microbiol. Biotechnol. 104, 9801–9822. 10.1007/s00253-020-10900-9

Seo, J.-A., Yu, J.-H., 2006. The phosducin-like protein PhnA is required for Gbetagamma-mediated signaling for vegetative growth, developmental control, and toxin biosynthesis in Aspergillus nidulans. Eukaryot. Cell 5, 400–410. 10.1128/EC.5.2.400-410.2006

Singer-Krüger, B., Jansen, R.-P., 2022. Proteomic Mapping by APEX2-Catalyzed Proximity Labeling in Saccharomyces cerevisiae Semipermeabilized Cells. Methods Mol. Biol. Clifton NJ 2477, 261–274. 10.1007/978-1-0716-2257-5_15

Todd, R.B., Fraser, J.A., Wong, K.H., Davis, M.A., Hynes, M.J., 2005. Nuclear Accumulation of the GATA Factor AreA in Response to Complete Nitrogen Starvation by Regulation of Nuclear Export. Eukaryot. Cell 4, 1646–1653. 10.1128/EC.4.10.1646-1653.2005

Xiong, Z., Lo, H.P., McMahon, K.-A., Parton, R.G., Hall, T.E., 2021. Proximity Dependent Biotin Labelling in Zebrafish for Proteome and Interactome Profiling. Bio-Protoc. 11, e4178. 10.21769/BioProtoc.4178

Yang, X., Wen, Z., Zhang, D., Li, Z., Li, D., Nagalakshmi, U., Dinesh-Kumar, S.P., Zhang, Y., 2021. Proximity labeling: an emerging tool for probing in planta molecular interactions. Plant Commun. 2, 100137. 10.1016/j.xplc.2020.100137

Zhang, Y., Li, Y., Yang, X., Wen, Z., Nagalakshmi, U., Dinesh-Kumar, S.P., 2020. TurboID-Based Proximity Labeling for In Planta Identification of Protein-Protein Interaction Networks. J. Vis. Exp. JoVE. 10.3791/60728

## References

Atanasoff-Kardjalieff, A. K., Studt, L., 2022. Secondary Metabolite Gene Regulation in Mycotoxigenic Fusarium Species: A Focus on Chromatin. Toxins. 14, 96.

Berger, H., et al., 2008. Dissecting individual steps of nitrogen transcription factor cooperation in the Aspergillus nidulans nitrate cluster. Mol Microbiol. 69, 1385–98.

Berger, H., et al., 2006. The GATA factor AreA regulates localization and in vivo binding site occupancy of the nitrate activator NirA. Mol Microbiol. 59, 433–46.

Bernreiter, A., et al., 2007. Nuclear export of the transcription factor NirA is a regulatory checkpoint for nitrate induction in Aspergillus nidulans. Molecular and Cellular Biology. 27, 791–802.

Boettcher, B., Barral, Y., 2013. The cell biology of open and closed mitosis. Nucleus. 4, 160–5.

Burger, G., et al., 1991. Molecular cloning and functional characterization of the pathway-specific regulatory gene nirA, which controls nitrate assimilation in Aspergillus nidulans. Mol Cell Biol. 11, 795–802.

Connolly, L. R., et al., 2013. The Fusarium graminearum histone H3 K27 methyltransferase KMT6 regulates development and expression of secondary metabolite gene clusters. PLoS genetics. 9, e1003916.

Cove, D. J., 1979. Genetic studies of nitrate assimilation in Aspergillus nidulans. Biol Rev Camb Philos Soc. 54, 291–327.

Davies, J. R., et al., 2004. Potential link between the NIMA mitotic kinase and nuclear membrane fission during mitotic exit in Aspergillus nidulans. Eukaryot Cell. 3, 1433–44.

Deng, L., et al., 2013. Dissociation kinetics of the streptavidin-biotin interaction measured using direct electrospray ionization mass spectrometry analysis. J Am Soc Mass Spectrom. 24, 49–56.

Gacek-Matthews, A., et al., 2016. KdmB, a Jumonji Histone H3 Demethylase, Regulates Genome-Wide H3K4 Trimethylation and Is Required for Normal Induction of Secondary Metabolism in Aspergillus nidulans. PLoS genetics. 12, e1006222.

Gacek, A., Strauss, J., 2012. The chromatin code of fungal secondary metabolite gene clusters. Appl Microbiol Biotechnol. 95, 1389–404.

Gallmetzer, A., et al., 2015. Reversible Oxidation of a Conserved Methionine in the Nuclear Export Sequence Determines Subcellular Distribution and Activity of the Fungal Nitrate Regulator NirA. PLoS Genetics. 11, e1005297.

Georgiou, X., et al., 2023. The interactome of the UapA transporter reveals putative new players in anterograde membrane cargo trafficking. Fungal Genetics and Biology. 169, 103840.

Harris, S. D., et al., 2009. Morphology and development in Aspergillus nidulans: a complex puzzle. Fungal Genet Biol. 46 Suppl 1, S82–S92.

Joseph, J. D., Means, A. R., 2000. Identification and characterization of two Ca2+/CaM-dependent protein kinases required for normal nuclear division in Aspergillus nidulans. J Biol Chem. 275, 38230–8.

Karahoda, B., et al., 2022. The KdmB-EcoA-RpdA-SntB chromatin complex binds regulatory genes and coordinates fungal development with mycotoxin synthesis. Nucleic Acids Research. 50, 9797–9813.

Kudla, B., et al., 1990. The regulatory gene areA mediating nitrogen metabolite repression in Aspergillus nidulans. Mutations affecting specificity of gene activation alter a loop residue of a putative zinc finger. Embo J. 9, 1355–64.

Lamb, H. K., et al., 2003. The negative transcriptional regulator NmrA discriminates between oxidized and reduced dinucleotides. The Journal of Biological Chemistry. 278, 32107–32114.

Langdon, T., et al., 1995. Mutational analysis reveals dispensability of the N-terminal region of the Aspergillus transcription factor mediating nitrogen metabolite repression. Mol Microbiol. 17, 877–88.

Lemmon, M. A., 2004. Pleckstrin homology domains: not just for phosphoinositides. Biochem Soc Trans. 32, 707–11.

Li, H., et al., 2021. Thiol-Cleavable Biotin for Chemical and Enzymatic Biotinylation and Its Application to Mitochondrial TurboID Proteomics. J Am Soc Mass Spectrom. 32, 2358–2365.

May, D. G., et al., 2020. Comparative Application of BioID and TurboID for Protein-Proximity Biotinylation. Cells. 9, 1070.

Muro-Pastor, M. I., et al., 1999. The GATA factor AreA is essential for chromatin remodelling in a eukaryotic bidirectional promoter. Embo J. 18, 1584–97.

Muro-Pastor, M. I., et al., 2004. A paradoxical mutant GATA factor. Eukaryot Cell. 3, 393–405.

Pfannenstiel, B. T., Keller, N. P., 2019. On top of biosynthetic gene clusters: How epigenetic machinery influences secondary metabolism in fungi. Biotechnol Adv. 37, 107345.

Punt, P. J., et al., 1995. The intergenic region between the divergently transcribed niiA and niaD genes of Aspergillus nidulans contains multiple NirA binding sites which act bidirectionally. Molecular and Cellular Biology. 15, 5688–5699.

Reyes-Dominguez, Y., et al., 2012. Heterochromatin influences the secondary metabolite profile in the plant pathogen Fusarium graminearum. Fungal Genet Biol. 49, 39–47.

Scazzocchio, C., 2000. The fungal GATA factors. Curr Opin Microbiol. 3, 126–31.

Scazzocchio, C., Arst, H. N., Regulation of nitrate assimilation in Aspergillus nidulans. In: J. L. Wray, J. R. Kinghorn, (Eds.), Molecular and genetic aspects of nitrate assimilation. Oxford Science Publications, Oxford, 1989, pp. 299–313.

Schinko, T., et al., 2010. Transcriptome analysis of nitrate assimilation in Aspergillus nidulans reveals connections to nitric oxide metabolism. Mol Microbiol. 78, 720–38.

Sosa Ponce, M. L., et al., 2024. Lipids and chromatin: a tale of intriguing connections shaping genomic landscapes. Trends Cell Biol.

Soyer, J. L., et al., 2014. Epigenetic control of effector gene expression in the plant pathogenic fungus Leptosphaeria maculans. PLoS Genet. 10, e1004227.

Stammers, D. K., et al., 2001. The structure of the negative transcriptional regulator NmrA reveals a structural superfamily which includes the short-chain dehydrogenase/reductases. Embo J. 20, 6619–26.

Strauss, J., et al., 1998. The regulator of nitrate assimilation in ascomycetes is a dimer which binds a nonrepeated, asymmetrical sequence. Molecular and Cellular Biology. 18, 1339–1348.

Studt, L., et al., 2016. Knock-down of the methyltransferase Kmt6 relieves H3K27me3 and results in induction of cryptic and otherwise silent secondary metabolite gene clusters in Fusarium fujikuroi. Environmental microbiology. 18, 4037–4054.

Trinkle-Mulcahy, L., 2019. Recent advances in proximity-based labeling methods for interactome mapping. F1000Res. 8.

Virag, A., Harris, S. D., 2006. Functional characterization of Aspergillus nidulans homologues of Saccharomyces cerevisiae Spa2 and Bud6. Eukaryot Cell. 5, 881–95.

Wong, K. H., et al., 2007. Transcriptional control of nmrA by the bZIP transcription factor MeaB reveals a new level of nitrogen regulation in Aspergillus nidulans. Mol Microbiol. 66, 534–51.

Yin, L. M., et al., 2020. Structural Characteristics, Binding Partners and Related Diseases of the Calponin Homology (CH) Domain. Front Cell Dev Biol. 8, 342.

